# Noradrenergic signaling in the rodent orbitofrontal cortex is required to update goal-directed actions

**DOI:** 10.1101/2022.06.30.498245

**Authors:** Juan-Carlos Cerpa, Alessandro Piccin, Margot Dehove, Marina Lavigne, Eric J. Kremer, Mathieu Wolff, Shauna L. Parkes, Etienne Coutureau

## Abstract

In a constantly changing environment, organisms must track the current relationship between actions and their specific consequences and use this information to guide decision-making. Such goal-directed behavior relies on circuits involving cortical and subcortical structures. Notably, a functional heterogeneity exists within the medial prefrontal, insular, and orbitofrontal cortices (OFC) in rodents. The role of the latter in goal-directed behavior has been debated, but recent data indicate that the ventral and lateral subregions of the OFC are needed to integrate changes in the relationships between actions and their outcomes. Neuromodulatory agents are also crucial components of prefrontal functions and behavioral flexibility might depend upon the noradrenergic modulation of prefrontal cortex. Therefore, we assessed whether noradrenergic innervation of the OFC plays a role in updating action-outcome relationships. We used an identity-based reversal task and found that depletion or chemogenetic silencing of noradrenergic inputs within the OFC rendered rats unable to associate new outcomes with previously acquired actions. Silencing of noradrenergic inputs in the medial prefrontal cortex or depletion of dopaminergic inputs in the OFC did not reproduce this deficit. Together, our results indicate that noradrenergic projections to the OFC are required to update goal-directed actions.

**GRAPHICAL ABSTRACT:** 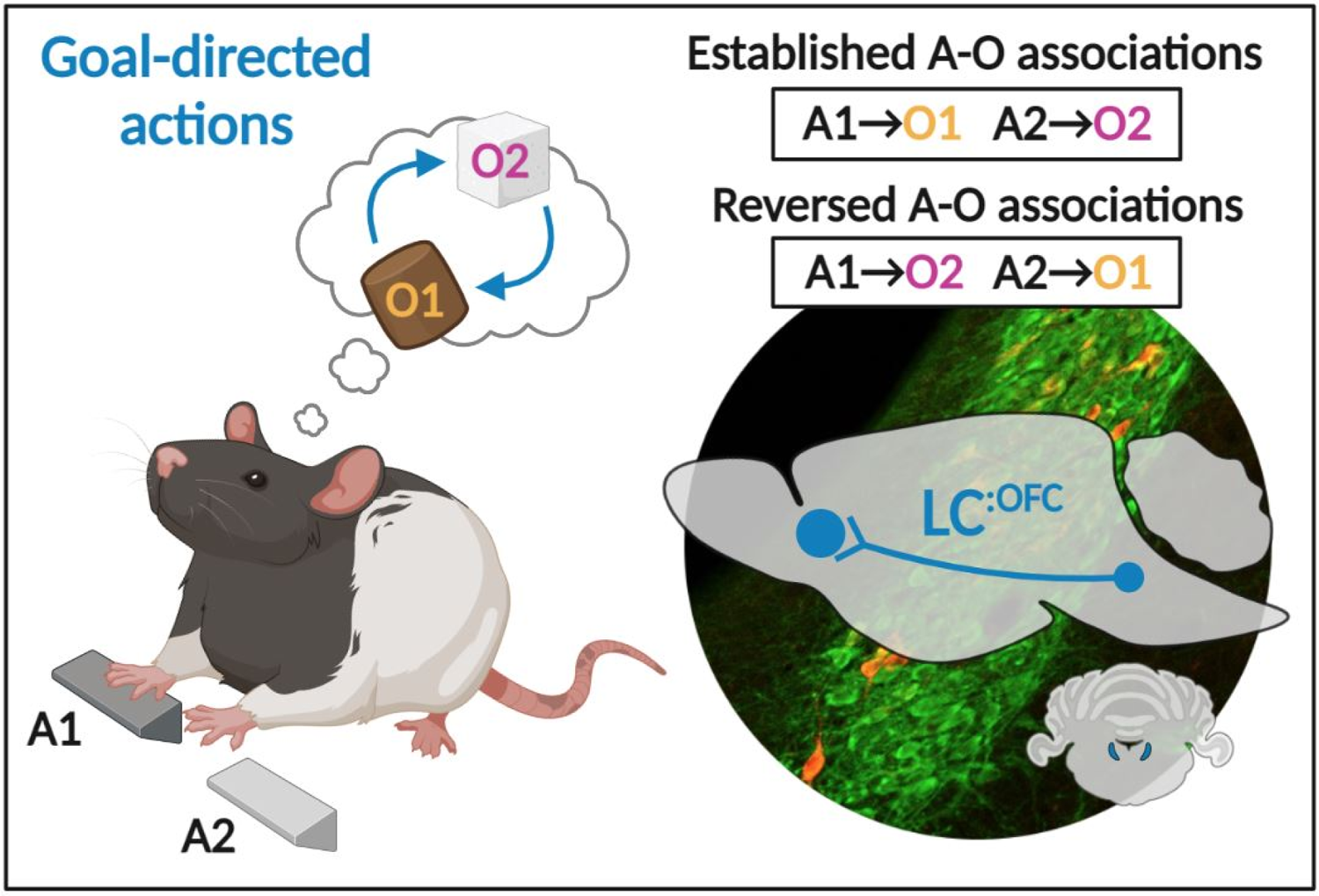

**HIGHLIGHTS:**

- Rats learn initial action-outcome associations in an instrumental task
- Noradrenergic depletion in the OFC prevents the encoding and expression of these associations following reversal learning
- Dopaminergic depletion in the OFC does not result in behavioral deficits
- LC:^OFC^ noradrenergic projections are required to update action-outcome associations

**IN BRIEF:** Cerpa et al. investigate whether noradrenergic projections from the locus coeruleus (LC) to the orbitofrontal cortex are involved in updating previously established goal-directed actions following environmental change. They find that these LC projections are required to both encode and express reversed action-outcome associations in rats.

## INTRODUCTION

Animals use their knowledge of an environment to engage in behaviors that meet their basic needs and desires. In a dynamic environment, an animal must also be able to update its understanding of the setting, particularly when the outcomes or consequences of its actions change. Numerous studies indicate that goal-directed behaviors are supported by the prefrontal cortex (PFC), and current research suggests a parcellation of functions within prefrontal regions in rodents (for recent reviews see Balleine, 2019, O’Doherty et al 2017, Coutureau & Parkes 2018). Specifically, the prelimbic region of the medial PFC (mPFC) is needed to initially acquire goal-directed actions and learn the relationship between distinct actions and their outcomes (Corbit & Balleine 2003, Killcross & Coutureau 2003, Tran-Tu-Yen et al 2009), whereas the gustatory region of the insular cortex is required to recall the current value of these outcomes to guide choice between competing actions (Balleine & Dickinson 2000, Parkes & Balleine 2013, Parkes et al 2015). In addition, the ventral (VO) and lateral (LO) subregions of the orbitofrontal cortex (OFC) are required to update previously established goal-directed actions (Parkes et al 2018). These data suggest that the VO and LO play critical roles in tracking the relationships between actions and their consequences.

Behavioral flexibility also requires activity in noradrenergic (NA) neurons of the locus coeruleus (LC), which are thought to track uncertainty in the current situation (Bouret & Sara 2004, Cope et al 2019, Jahn et al 2018, McGaughy et al 2008, Tait et al 2007, Tervo et al 2014, Uematsu et al 2017). Most notably, compelling theoretical models hypothesize that the LC interacts with the PFC to support behavioral flexibility (Sara & Bouret 2012). Taken together, these data raise the intriguing possibility that LC NA innervation of the OFC might be needed to update previously established goal-directed actions (Chandler et al 2013, Agster et al 2013, Sadacca et al 2016, Cerpa et al 2019, Cerpa et al 2021). To investigate this possibility, we examined whether LC NA inputs to the OFC are required to learn that a specific outcome associated with a given action has changed, and to recall this information to guide choice.

Specifically, after learning initial action-outcome (A-O) associations, rats were required to flexibly encode and use new associations during an instrumental reversal task. First, we depleted NA fibers in the OFC using anti-DβH-saporin and observed a profound deficit in the ability to use the reversed A-O associations to guide choice. We then found that this deficit is not present when we depleted dopaminergic signaling in the OFC. Finally, we investigated the temporal and anatomical specificity of this effect using viral vector-mediated expression of an inhibitory DREADD. We found that silencing LC:^OFC^, but not LC:^mPFC^, projections impaired the ability to acquire and express the reversed instrumental contingencies. Collectively, these data demonstrate that LC NA projections to the OFC are required for both encoding and recalling the identity of an expected instrumental outcome, specifically when that identity has changed.

## RESULTS

### Initial goal-directed learning does not require NA signaling in the OFC

We first assessed if the initial acquisition and expression of goal-directed actions requires NA signaling in the OFC using the behavioural design (**Figure 1A**). Rats in the Pre group were injected with the anti-DβH saporin toxin (SAP) before the initial instrumental training, during which responding on one action (A1) earned O1 (sucrose or grain pellets, counterbalanced), and responding on the other (A2) earned O2 (grain or sucrose pellets, counterbalanced). Rats in group Post were similarly trained, but were injected with SAP following this initial stage. To deplete NA projections, rats were given bilateral injections of SAP or inactive anti-IgG saporin (CTL) targeting the VO and LO. SAP infusion resulted in extensive NA fiber loss in the VO and LO, as revealed by a main effect of group (F_(1,53)_ = 276.95, p <0.001) and region (F_(1,53)_ = 12.31, p <0.01), and a significant group (CTL *vs* SAP) X region (VO *vs* LO) interaction effect was also detected (F_(1,53)_=12.42,, p <0.01) (**Figure 1B**). Simple effect analyses confirmed a significant reduction of NA fibers density in both the VO (F_(1,53)_ = 282.27, p <0.001) and the LO (F_(1,53)_ = 232.23, p <0.001) following SAP injection. By contrast, we found a higher volume of NA fibers in the VO of the control group, as compared to the LO (F_(1,53)_ = 23.45, p <0.001), a result consistent with previous findings (Cerpa et al 2019). However, unexpectedly, significant fiber loss was also observed in the medial prefrontal cortex (mPFC; F_(1,53)_ = 195.57, p <0.001; **Figure 1C**).

**Figure 1.**
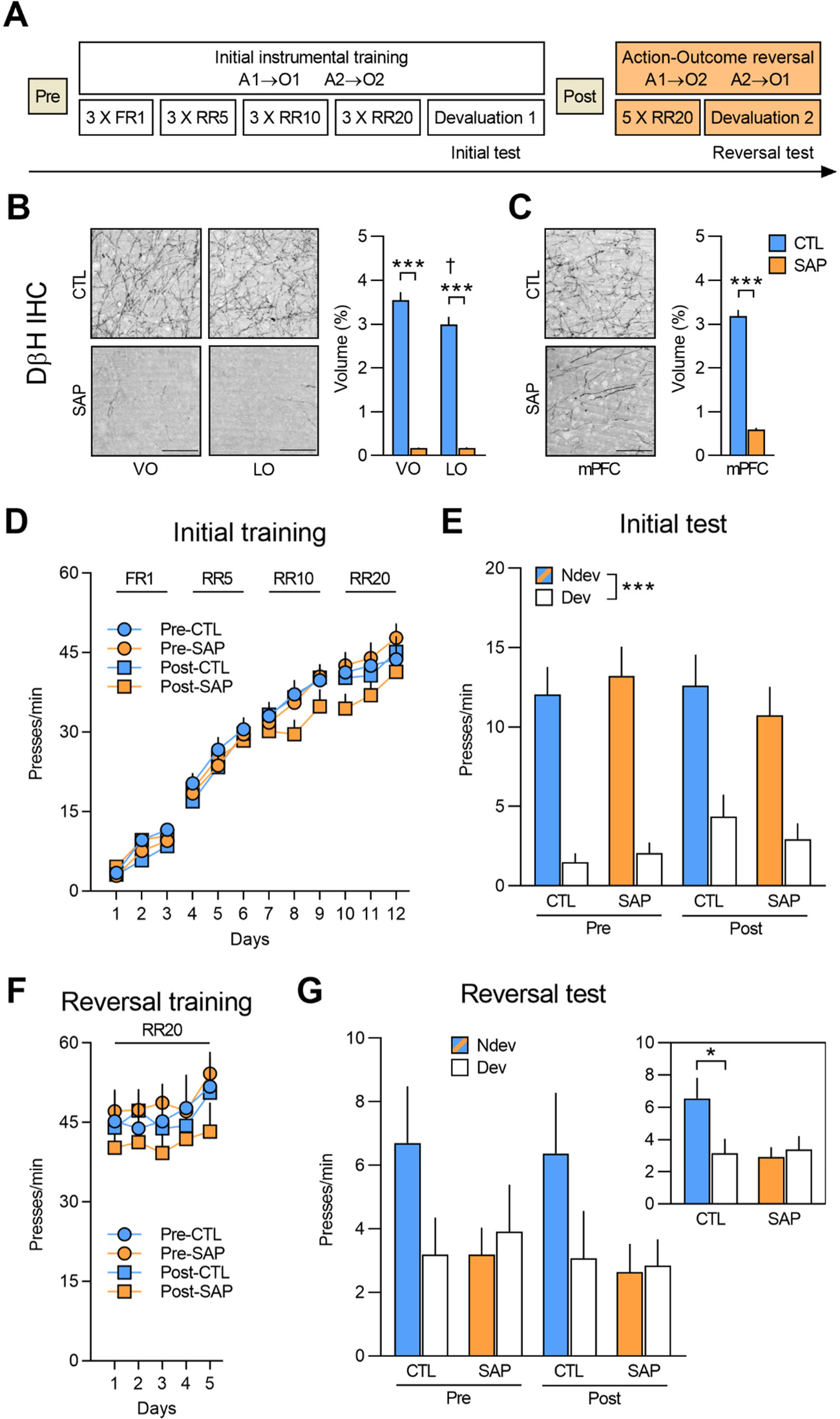
(**A**) Experimental timeline for rats injected with anti-DβH saporin in the OFC before (Pre) and after (Post) initial instrumental training and outcome devaluation (Pre-CTL n = 14, Pre-SAP n = 15, Post-CTL n = 13, Post-SAP n = 15). (**B**) Representative microphotographs of noradrenergic depletion and DβH fibre volume in the VO (+3.7 mm from Bregma) and LO (+3.7 mm from Bregma) following toxin injection. (**C**) Representative microphotograph of noradrenergic depletion and DβH fibre volume in the mPFC (medial orbitofrontal+A32d+A32v; +4.4 mm from Bregma) following toxin injection. (**D**) Rate of lever pressing across initial training (A1-O1; A2-O2), collapsed across the two actions. (**E**) Initial instrumental test in extinction following satiety-induced devaluation. (**F**) Rate of lever pressing across reversal training (A1-O2; A2-O1), collapsed across the two actions. (**G**) Reversal instrumental test in extinction following satiety-induced devaluation. The inlet shows data grouped for CTL (Pre and Post) and SAP groups (Pre and Post). Data are presented as mean + S.E.M. *p <0.05, ***p <0.001, ^†^p <0.05 LO *vs* VO CTL group. Scale bars: 100 μm.

We found that all rats acquired the lever pressing response, with their rate of lever pressing increasing across days (F_(1,53)_ = 508.30, p <0.001). No differences were found between Pre versus Post groups (F_(1,53)_ = 0.96, p = 0.33) or between CTL and SAP groups (F_(1,53)_ = 0.287, p = 0.59) and there were no significant interactions (all F_(1,53)_ values <1.18, p values > 0.28) (**Figure 1D**). All groups also showed sensitivity to the change in value during the outcome devaluation test, indicating that rats learned the action-outcome associations and the current value of the outcomes value; i.e., goal-directed behaviour was intact (**Figure 1E**). Indeed, we found a significant effect of devaluation (Ndev *vs* Dev; F_(1,53)_=79.62, p <0.001) but no effect of group (Pre *vs* Post; F_(1,53)_=0.213, p=0.65) or treatment (CTL *vs* SAP; F_(1,53)_=0.152, p=0.70), and no significant interactions between these factors (all F_(1,53)_ values <1.78, p values >0.18). In addition, when given concurrent access to both outcomes, all groups consumed more of the non-devalued outcome, thereby substantiating the efficacy of the satiety-induced outcome devaluation procedure (**Supp. Figure 1A**). Thus, NA depletion in the OFC and mPFC does not affect the initial learning or expression of goal-directed actions.

### NA signaling in the OFC is required to adapt to changes in outcome identity

Next, we tested whether NA depletion affected the ability to encode and express new action-outcome associations using an instrumental outcome-identity reversal paradigm. We found that performance increased across reversal training days (F_(1,53)_ = 22.73, p <0.001), but there was no effect of group (F_(1,53)_ = 0.80, p = 0.38), treatment (F_(1,53)_ = 0.08, p=0.78), or any significant interactions between these factors (all F_(1,53)_ values <1.86, p values >0.17) (**Figure 1F**). We then evaluated if the rats were able to use the reversed associations in the outcome devaluation test (**Figure 1G**). While the control groups reduced lever pressing associated with the devalued outcome, NA-depleted animals did not. Statistical analyses confirmed this observation revealing no main effects of group, treatment, or devaluation (all F_(1,53)_ values <2.88, p values >0.10), and no significant devaluation X group interaction (F_(1,53)_ = 0.007, p = 0.93). However, there was a significant devaluation X treatment interaction (F_(1,53)_ = 5.00, p <0.05) and simple effect analyses confirmed that the control groups biased their responding toward the lever associated with the non-devalued outcome (F_(1,53)_ = 7.35, p <0.01). Notably, rats in the depleted groups did not (F_(1,53)_ = 0.15, p = 0.70). Importantly, all groups rejected the devalued food during the consumption test (**Supp. Figure 1B**). These results show that depletion of NA innervation to OFC and mPFC renders rats unable to associate new outcomes to acquired actions. Importantly, this deficit was present in rats that received NA depletion before (group Pre) and after (group Post) learning the initial action-outcome associations, which is consistent with a process of updating established goal-directed actions.

### DA signaling in the OFC is not required to adapt to changes in outcome identity

Next, we examined whether dopaminergic (DA) signaling in the OFC is required for updating goal-directed actions. We used the behavioral procedure previously described (see **Figure 1A**) and all rats underwent surgery following initial acquisition and expression of goal-directed behaviour (see **Suppl. Figure 2A-C, Suppl. Figure 3A-C** for behavioral results from this initial phase). We first assessed the impact of full catecholaminergic (CA) deletion on outcome-identity reversal by infusing the CA-targeting toxin 6-OHDA in the OFC (6-OHDA n = 12; control n = 8). We then depleted DA afferents by combining the 6-OHDA infusion with a systemic injection of desipramine (Desi) to protect NA fibers (6-OHDA + Desi, n = 9; control n = 8). In the OFC, DA depletion was effective as assessed by tyrosine hydroxylase (TH) immunoreactivity (**Figure 2A**). We found a a significant within-subject effect of region (VO versus LO; F_(1,26)_ = 8.18, p <0.01) and fewer TH-positive fibres in group 6-OHDA (n = 12) compared to groups control CTL (n = 8) and 6-OHDA + Desi (n = 9) (F_(1,26)_ = 41.14, p <0.001), which also differed (F_(1,26)_ = 15.7, p <0.01). There was also a significant interaction between these factors (F_(1,26)_ = 15.17, p <0.01) and simple effects revealed that the 6-OHDA + Desi cohort had fewer immunoreactive TH fibres in the LO versus the VO (F_(1,26)_ = 20.16, p <0.001). We found no significant difference between LO and VO in the controls (CTL) (F_(1,26)_ = 1.14, p = 0.30) or 6-OHDA (F_(1,26)_ = 2.74, p = 0.11). Importantly, significantly fewer TH-positive fibres were also present in the mPFC (MO + A32d + A32v) for the 6-OHDA (F_(1,26)_ = 10.78, p <0.01), but not 6OHDA+desi cohorts (F_(1,26)_ = 2.07, p = 0.16) (**Figure 2B**), indicating that, as in the previous experiment, depletions were not restricted to the OFC.

**Figure 2.**
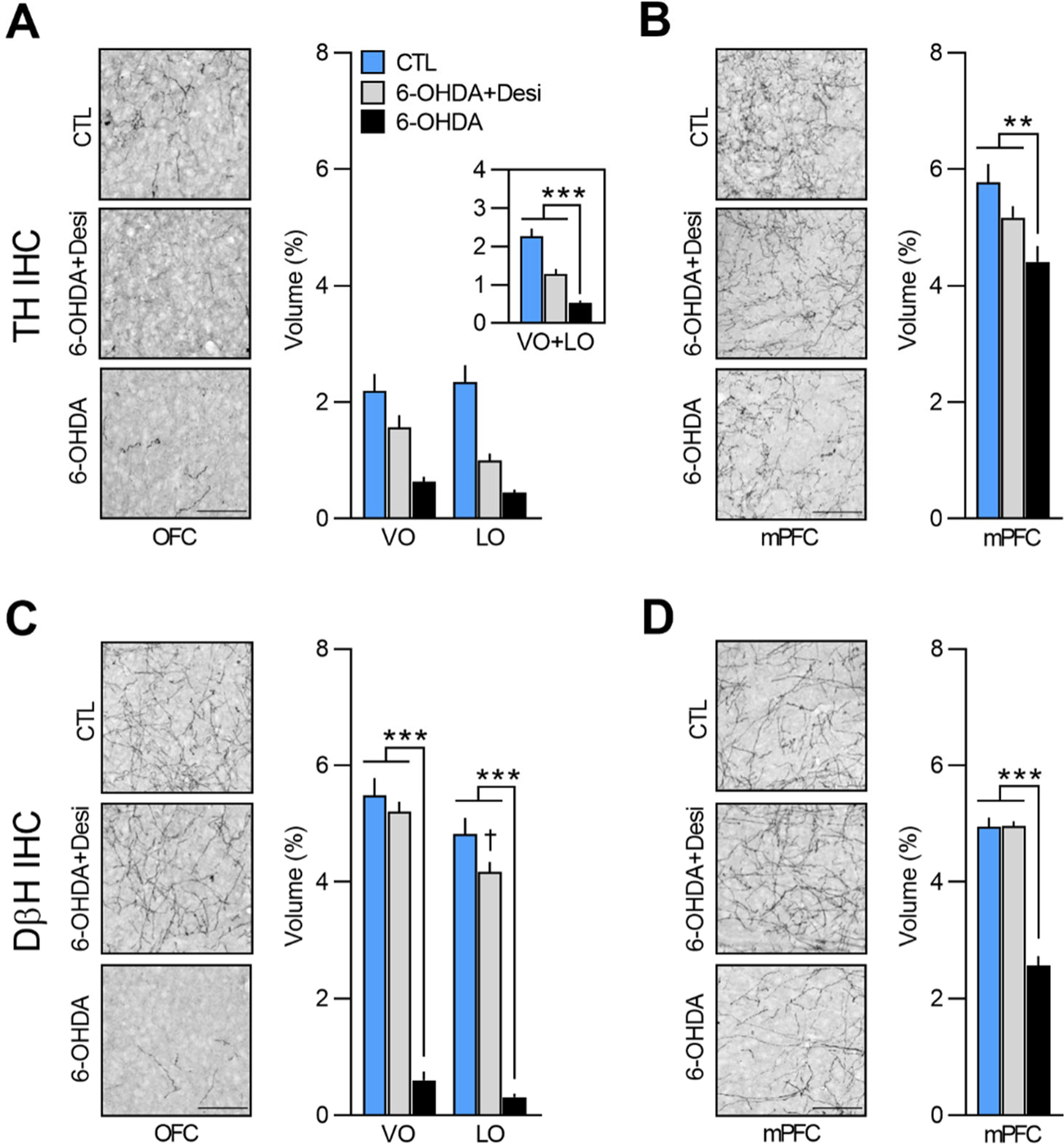
(**A**, **B**) Representative microphotographs and percentage of dopaminergic depletion in the OFC (VO and LO, +3.7 mm from Bregma) and in the mPFC (MO, A32d and A32v, +4.4 mm from Bregma) following toxin injection. (**C**, **D**) Representative microphotographs and percentage of noradrenergic depletion in the OFC (VO and LO, +3.7 mm from Bregma) and in the mPFC (MO, A32d and A32v, +4.4 mm from Bregma) following toxin injection. Data are presented as mean + S.E.M. **p <0.01, ***p <0.001, ^✝^p <0.05 *vs* CTL group. Scale bars: 100 μm.

The volume of DβH+ fibres in OFC and mPFC also differed. In the OFC, main effects analyses revealed a significant within-subject effect of region (VO versus LO; F_(1,26)_ = 53.24, p <0.001) and fewer TH+ fibres in the 6-OHDA group (n = 12) compared to CTL (n = 8) and 6-OHDA+Desi (n = 9) (F_(1,26)_ = 511.49, p <0.001), which did not differ (F_(1,26)_ = 3.24, p = 0.08) (**Figure 2C**). There was also a region x treatment interaction (F_(1,26)_ = 9.52, p <0.01) and simple effect analyses revealed that 6-OHDA treatment induced a significant reduction in the volume of NA fibers in both the VO (F_(1,26)_ = 400.70, p <0.001) and the LO (F_(1,26)_ = 459.47, p <0.001). Significantly fewer DβH+ fibers were also observed for the 6OHDA+Desi group in the LO (F_(1,26)_ = 6.42, p=0.02), but not in the VO group (F_(1,26)_ = 0.83, p=0.37). In the mPFC, a significant reduction in NA innervation was observed in the 6-OHDA group (F_(1,26)_ = 65.49, p <0.001), but not the 6OHDA+Desi group (F_(1,26)_ = 0.55, p=0.47), an effect that was previously observed (Figure 1C) and further indicates that our depletion was not specific to OFC (**Figure 2D**).

Following surgery and recovery, animals were trained with reversed instrumental associations. We found that rats with 6-OHDA infusions responded more during reversal training than the controls (F_(1,18)_ = 14.7, p <0.01) (**Figure 3B**). There was also an overall increase in response rate (F_(1,18)_ = 44.57, p <0.001) and a significant group x day interaction (F_(1,18)_ = 7.24, p <0.05), indicating that this increase in responding was greater for the 6-OHDA group than for controls. However, during the outcome devaluation test following reversal training (**Figure 3C**), control rats successfully adjusted their behaviour according to the new action-outcome contingencies, but 6-OHDA group did not. Statistical analysis revealed no main effect of group (F_(1,18)_ < 0.001, p >0.975), but a main effect of devaluation (F_(1,18)_ = 8.72, p <0.01) and a group x devaluation interaction that approached statistical significance (F_(1,18)_=3.90, p=0.06). Simple effect analyses confirmed that control rats responded more on the lever associated with the devalued outcome (F_(1,18)_ = 10.12, p <0.01), while 6-OHDA rats did not show this preference (F_(1,18)_ = 0.6, p = 0.45). Importantly, when given access to both rewards, all groups consumed more of the non-devalued food than the devalued food (**Suppl. Figure 2D**).

**Figure 3.**
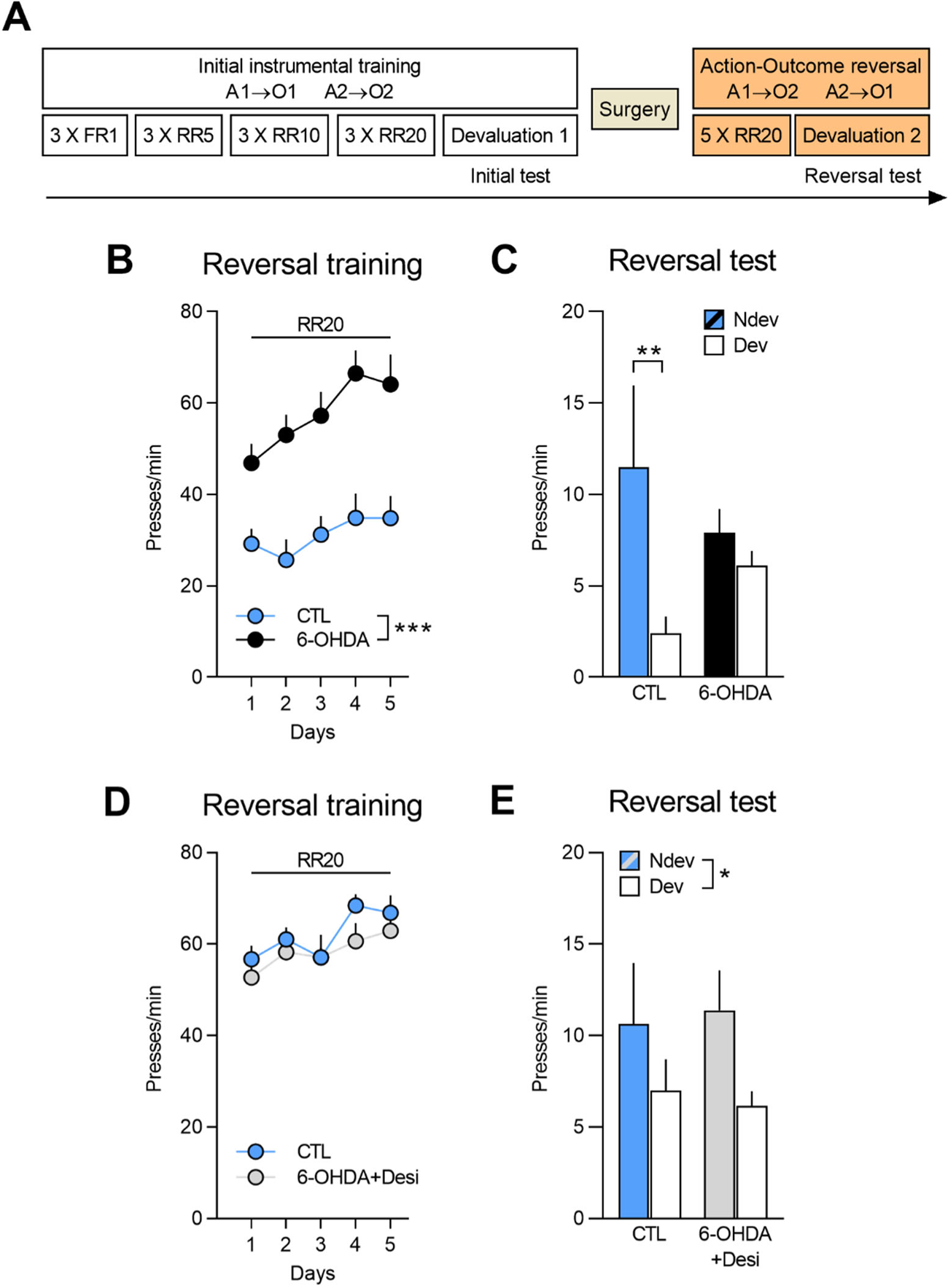
(**A**) Experimental timeline. After the initial instrumental training and outcome devaluation testing, rats were injected in the OFC with either vehicle (CTL n = 8; CTL n = 8), 6-OHDA coupled with desipramine (to specifically target DA neurons, 6-OHDA+Desi n = 9) or 6-OHDA alone (to target all CA neurons, n = 8). (**B**, **D**) Rate of lever pressing across reversal training, data is presented collapsed across the two actions. (**C**, **E**) Reversal instrumental test in extinction following satiety-induced devaluation. Data are presented as mean + S.E.M. *p <0.05, **p <0.01, ***p <0.001.

**Figure 3D** shows responding during reversal training for 6-OHDA + Desi and control groups. Lever pressing increased across reversal training days (F_(1,15)_ = 15.79, p <0.01) and there was no main effect of group (F_(1,15)_ = 0.52, p = 0.48) or group x day interaction (F_(1,15)_ = 0.157, p = 0.70). Moreover, we found that both groups showed goal-directed behaviour and biased their choice towards the action associated with the non-devalued outcome (Figure 3E). Statistical analyses confirmed a main effect of devaluation (F_(1,15)_=5.25, p <0.05), but no main effect of group (F_(1,15)_ < 0.001, p=0.975) or group x devaluation interaction (F_(1,15)_=0.168, p=0.69). Notably, both groups consumed more of the non-devalued food during the consumption test (**Suppl. Fig 3D**).

These results further support a specific role of NA in updating previously learned goal-directed actions. We show that full CA depletion (DA+NA) in the OFC and mPFC impairs performance in an outcome-identity reversal task, while depletion restricted to DA innervation leaves performance intact.

### Selective expression of inhibitory DREADDs in LC:^OFC^ or LC:^mPFC^ NA projections

In the previous two approaches, we used pharmacologic ablation to target NA signaling in the OFC. However, injection of anti-DHβ-saporin or 6-OHDA in the OFC also caused a significant reduction of fibres in the mPFC, most likely because of NA projections connecting the two cortical areas and/or fibers crossing the OFC before entering the mPFC (Chandler & Waterhouse 2012, 2013, 2014). As such, while we were able to demonstrate that NA but not DA signaling in the prefrontal cortex is necessary to adapt to changes in outcome identity, we could not conclusively attribute our behavioural effects to NA depletion in the OFC. Moreover, given that our approach involved permanent lesions of NA fibres, we were unable to ascertain if NA signaling was required to encode and/or recall the new action-outcome associations.

Therefore, to address the regional and temporal specificity of the behavioural effect, we generated CAV2-PRS-hM4D-hSyn-mCherry, a canine adenovirus vector containing PRS, a noradrenergic-specific promoter, driving an HA-tagged hM4D, an inhibitory DREADD, and an mCherry expression cassette. CAV-2 vectors are readily taken up at presynapse and trafficked, via retrograde transport to the soma of projecting neurons CAV2-PRS-hM4D-hSyn-mCherry was infused in either the OFC or the mPFC to targeted either LC:^OFC^ or LC:^mPFC^ NA projections (**Figure 4A**).**Figure 4B** shows retrograde transport of the vector and mCherry in NA cells of the LC following injection of CAV2-PRS-hM4D-hSyn-mCherry in the OFC. The colocalization of mCherry and HA immunoreactivity in the LC indicates a selective expression of HA-hM4D. As expected, while mCherry staining is present at injection sites, reflecting local cortico-cortical connections that are not NA dependent, HA-immunoreactive cell bodies were found exclusively in the LC. These data are consistent with NA-specific expression of the HA-tagged Hm4D due to PRS, and nonselective expression of mCherry expression which is under the control of hSyn, driving pan-neuronal expression (**Figure 4D**; **Suppl. Figure 4C**).

**Figure 4.**
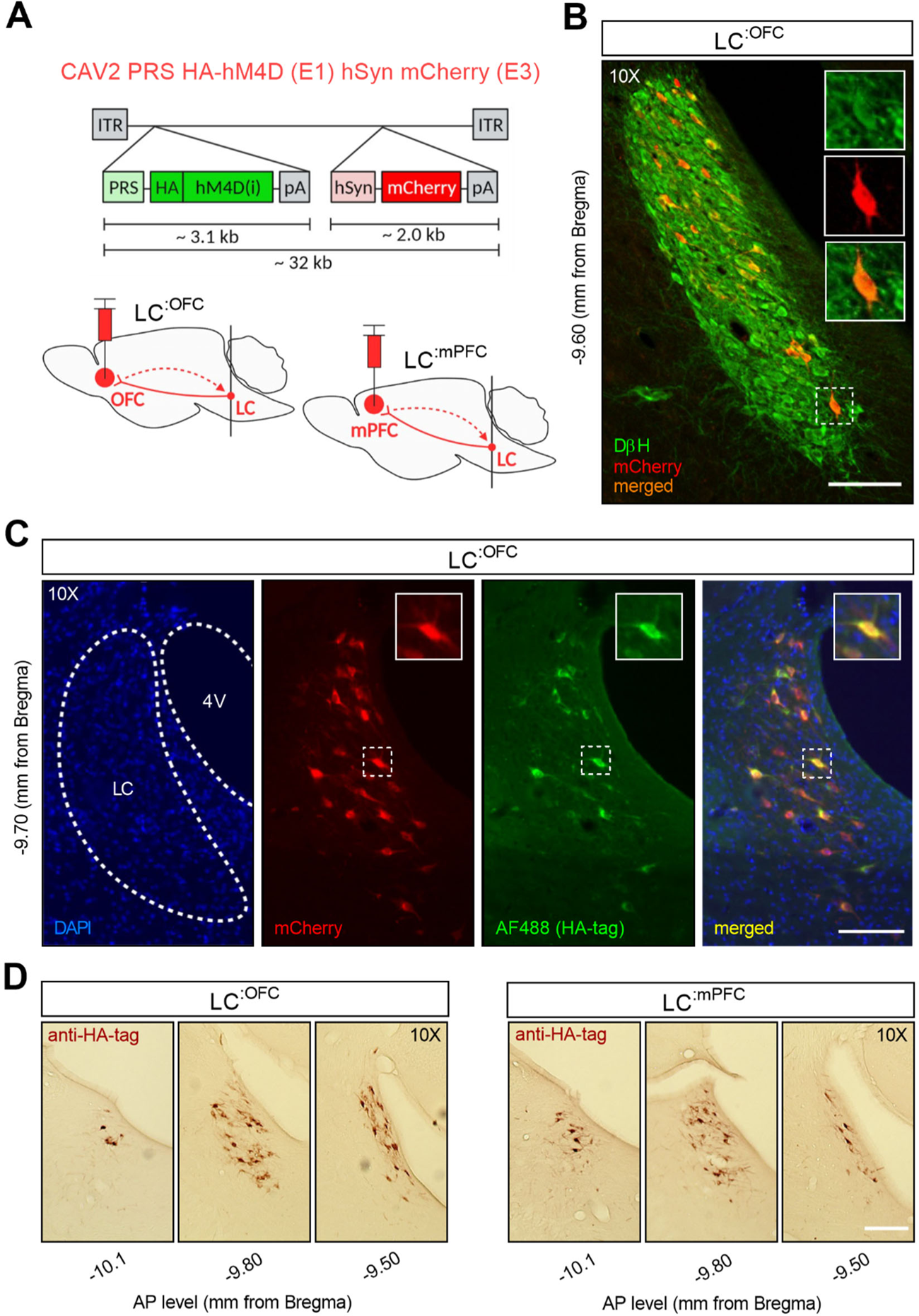
(**A**) CAV2-PRS-hM4D-mCherry, a vector bearing a noradrenergic-specific promoter (PRS) an inhibitory DREADD tagged with HA (hM4Di), an mCherry expression cassette and sites of injection. (**B**) Immunofluorescent staining for DβH and mCherry in the LC of a representative rat injected with CAV2-PRS-hM4D-hSyn-mCherry in the OFC. (**C**) High colocalization of immunofluorescent staining for HA (tag of inhibitory DREADDs) and mCherry in the LC of the same representative rat injected in the OFC. (**D**) Comparison of antero-posterior DAB staining for HA in two representative rats, one injected in the OFC, the other in the mPFC. Scale bars: 100 μm.

### Silencing of LC:^OFC^, but not LC:^mPFC^, projections impairs adaptation to changes in the action-outcome association

Rats were trained and tested as shown in **Figure 5A**. Following initial instrumental training and outcome devaluation testing (**Suppl. Figure 5A-C** for LC:^OFC^ and **Suppl. Figure 6A-C** for LC:^mPFC^), they were divided in two groups according to their initial performance: one group that would receive vehicle (Veh, -) during reversal training and one group that would receive deschloroclozapine dihydrochloride (DCZ, +), a high affinity and selective agonist for hM4D (Nagai et al., 2020; Nentwig et al., 2021; Oyama et al., 2022).

**Figure 5.**
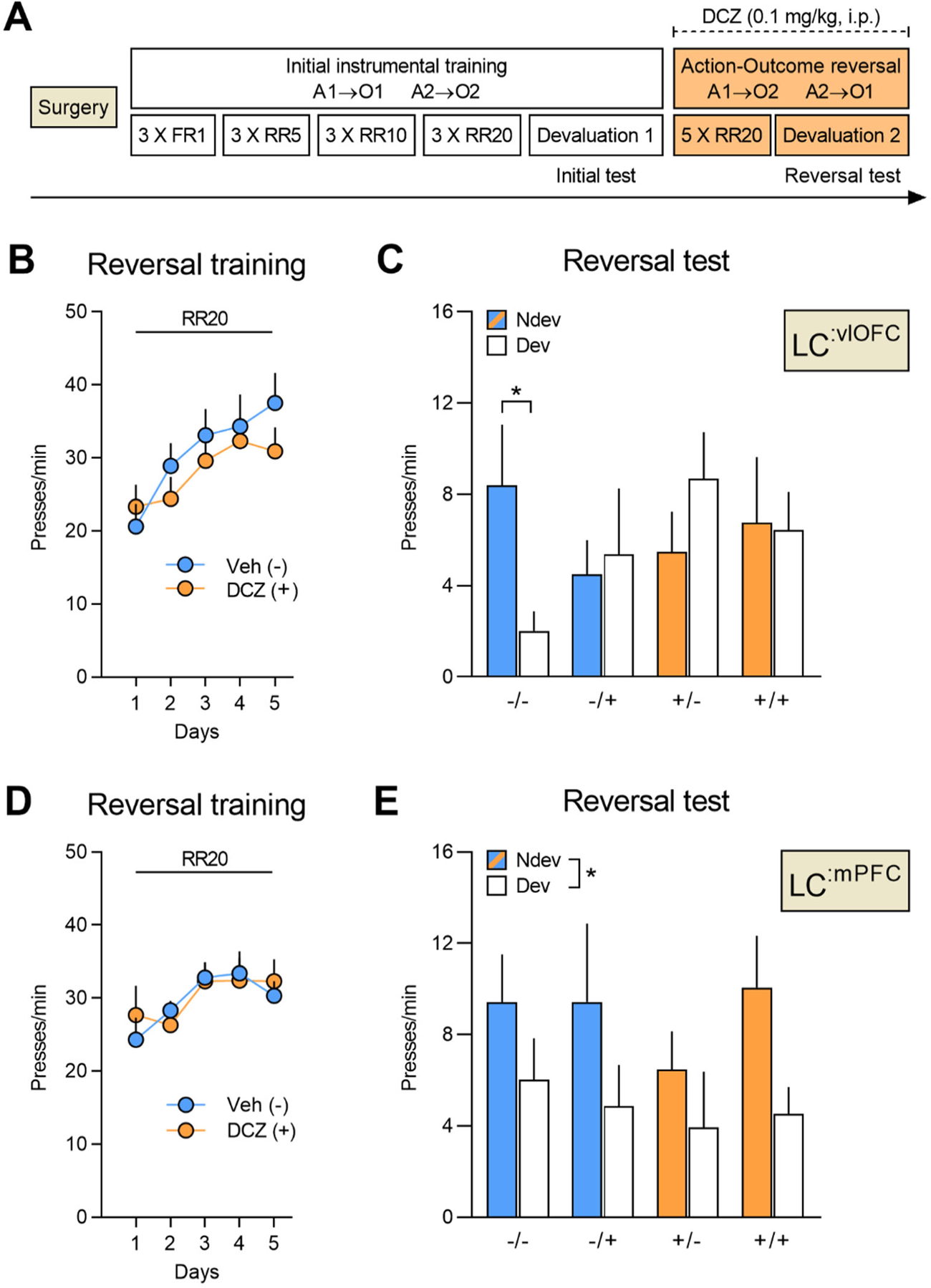
(**A**) Timeline for rats injected with CAV2-PRS-hM4D-hSyn-mCherry in either the OFC or the mPFC following reversal training. Each rat was injected with either vehicle (-) or DCZ (+) during reversal training and was then tested twice, once under DCZ and once under vehicle with the test order counterbalanced. (**B**) Reversal training in rats injected in the OFC (Veh n = 12; DCZ n = 13), data is presented collapsed across the two actions (A1-O2; A2-O1). (**C**) Reversal instrumental test following satiety-induced devaluation in rats injected in the OFC. (**D**) Reversal training in rats injected in the mPFC (Veh n = 8; DCZ n = 9), data is presented collapsed across the two actions (A1-O2; A2-O1). (**E**) Reversal instrumental test following satiety-induced devaluation in rats injected in the mPFC. Data are presented as mean + S.E.M. *p <0.05.

A linear trend across reversal training sessions was detected for OFC-injected rats (F_(1,23)_ = 37.44, p <0.001), but not mPFC-injected rats (F_(1,15)_ = 0.16, p = 0.69), with no difference between the Veh and the DCZ group for OFC-(F_(1,23)_ = 0.39, p = 0.54) or mPFC-injected rats (F_(1,15)_ = 0.01, p = 0.92), and no interactions between these factors (largest F_(1,23)_ value = 2.82, p = 0.11) ( **Figure 5B** & **D**). All rats then underwent two outcome devaluation tests, once under vehicle injection (-) and once under DCZ injection (+), with the order counterbalanced. This yielded a 2 (between) x 2 (within) factorial design with 4 conditions of interest: vehicle during training and test (-/-), vehicle during training and DCZ during test (-/+), DCZ during training and vehicle at test (+/-), and DCZ during training and test (+/+).

We found that only rats in the control group (-/-) showed goal-directed behaviour and performed the action associated with the non-devalued outcome more than the action associated with the devalued outcome (**Figure 5C**). By contrast, rats with bilateral LC:^OFC^ NA projections silenced during the reversal training (+/- and +/+) or the outcome devaluation test (-/+ and +/+) failed to display this preference in responding. The between- and within-subject main effects were not significant (largest F_(1,46)_ value = 1.22, p = 0.28), but there was a significant three-way interaction (devaluation X treatment during acquisition X treatment during test: F_(1,23)_ = 5.45, p <0.05). Simple effects analyses showed that only rats that received vehicle during both training and test (-/-) biased their choice toward the lever associated with the non-devalued outcome (F_(1,23)_ = 7.10, p <0.05), while the -/+ (F_(1,23)_=0.09, p=0.77), +/- (F_(1,23)_= 1.22, p=0.28), and +/+ groups (F_(1,23)_=0.01, p=0.92) did not. By contrast, silencing LC:^mPFC^ NA projections left goal-directed behaviour intact (main effect of devaluation: F_(1,15)_=5.763, p <0.05). We found no main effect of treatment during acquisition (F_(1,15)_=0.35, p=0.56) or treatment during test (F_(1,15)_=0.53, p=0.47) and no significant interactions between these factors (largest F_(1,15)_ value=1.62, p values >0.21) (**Figure 5E**).

Importantly, consumption tests performed immediately after the reversal tests revealed that all groups consumed more of the non-devalued outcome indicating that the satiety-induced devaluation was effective and that DCZ injections did not disrupt the rats’ ability to distinguish between devalued and non-devalued rewards (**Suppl. Figure 5D** and **6D** for LC:^OFC^ and LC:^mPFC^, respectively). Together, these results led us to conclude that LC NA projections to the OFC, but not to the mPFC, are required to both encode and recall changes in the identity of the expected outcome.

## DISCUSSION

Goal-directed actions are the expression of learned associations between an action and the outcome it produces. These associations are however flexible, being amenable to updating when the identity of the outcome changes. Our data demonstrate that NA inputs to the orbitofrontal cortex are specifically required for this updating process. This conclusion is based on a body of complementary data. First, we demonstrated that animals with a loss of NA inputs in the OFC can initially learn and express action–outcome contingencies, but are impaired when the identity of the outcome has been modified. Importantly, such deficits were also observed when NA depletion occurred immediately before the encoding of the new action-outcomes contingencies. We then showed that this impairment was selective to NA inputs to the OFC, because combined depletions of DA and NA, but not of DA alone, induced a profound deficit in outcome reversal. Finally, we investigated the temporal and anatomical specificity of this effect using a selective NA retrograde virus carrying inhibitory DREADDs to selectively target either LC:^OFC^ or LC:^mPFC^ pathways. We found that silencing LC:^OFC^, but not LC:^mPFC^, projections impaired the rats’ ability to acquire and express the reversed instrumental contingencies. Taken together, these results reveal the NA regulation of goal-directed behaviour.

### NA inputs into the OFC, but not the mPFC, are required for action-outcome updating

Depletion of NA inputs was achieved using neurochemical toxins (anti-DβH saporin or 6-OHDA). Both toxins led to a dramatic decrease in NA fiber density in the ventral and lateral OFC, and in the medial prefrontal cortex. This pattern is unlikely due to an overlap between LC projections to the mPFC and the OFC, given that divergent prefrontal projections from the LC exist but are limited (Chandler et al. 2013, Foote et al. 1983, Fuxe et al. 1968, Levitt & Moore 1978, Lewis & Morrison 1989, Morrison et al. 1978, Uematsu et al. 2017). It is therefore more likely that the depletion in the mPFC results from the crossing of the OFC by NA fibers *en route* to the mPFC, as recently suggested (Cerpa et al. 2019). In the present study, a high degree of anatomical selectivity was achieved by using a CAV-2 vector carrying the noradrenergic promoter PRS to target either the LC:^mPFC^ or the LC:^OFC^ pathways (Hayat et al; 2020; Hirschberg et al., 2017).

A key finding is that NA inputs to the OFC are specifically required for updating the association between an action and its outcome. Indeed, a similar impairment in goal-directed behaviour resulted from NA depletion performed either prior to initial training or prior to reversal training, which indicates that the reversal period is critically reliant on NA inputs. In addition, chemogenetic silencing of the LC:^OFC^ pathway before the reversal training, or before testing also produced similar impairments, which further demonstrates that OFC NA inputs are required for both the encoding and the recall of new action-outcome associations. These results are consistent with recent views on the role of the OFC in goal-directed behaviour (Parkes et al., 2018; Panayi & Killcross, 2018; Cerpa et al., 2021).

In contrast to NA inputs to the OFC, our results show that NA input to the mPFC is not required for responding based on initial or reversed instrumental contingencies. These data add to the current literature indicating a major dissociation in the role of NA inputs to different prefrontal regions (Robbins & Arnsten 2009). Indeed, NA input to mPFC is required for attentional regulation since lesioning NA inputs (McGaughy et al 2008, Newman et al 2008), or chemogenetic inhibition of NA-mPFC (Cope et al 2019) alter attentional set-shifting, while NA recapture inhibition via atomoxetine improves it (Newman et al 2008). Similar systematic investigation is not yet available but OFC NA depletion can alter cue-outcome reversal, but not dimensional shift, in an attentional set-shifting task (Mokler et al 2017).

### NA but not DA inputs to the OFC are required for action-outcome updating

Using a strategy which allows for a differential depletion of DA and/or NA fibers, we found that NA-dependent mechanisms are required during the encoding and recall of new action-outcome. The role of cortical DA-dependent mechanisms in goal-directed behaviour remains poorly understood. However, we previously demonstrated that dopaminergic signaling in the mPFC plays a critical role in the detection of contingency degradation (Naneix et al 2009, 2012, 2013). Such detection is likely to involve the processing of non-expected rewards which induces, at the level of mPFC, a DA-dependent reward prediction error signal (Montague et al 2004, Schultz & Dickinson 2000). These results therefore raise the intriguing possibility that the coordination of goal-directed behaviour under environmental changes might depend on a DA-mPFC system to adapt to causal contingencies and NA-OFC system to adapt to changes in outcome identity (Cerpa et al., 2021).

### Updating goal-directed behaviour

When trained on reversed contingencies, animals encode the new action-outcome associations (Fresno et al 2019, Parkes et al 2018). Under similar experimental conditions, past research has shown that reversal learning performance is the result of updating prior existing action-outcome contingencies without unlearning the initial ones (Bradfield & Balleine 2017). In other words, the animals build a partition between a state for the new contingencies and the initial state of old contingencies (Hart & Bradfield, 2020). Current research (McDannald et al 2011, Sadacca et al 2016, Wikenheiser & Schoenbaum 2016) has proposed that the OFC is critically involved in the partition of information when task states change without explicit notice (Wilson et al 2014). Consistent with this view, chemogenetic inhibition of the OFC (ventral and lateral) impairs goal-directed responding following identity reversal (Parkes et al 2018; Howard & Kahnt, 2021). Here, we found a similar deficit following lesion of NA inputs to the OFC. Given that the deficit in goal-directed behavior was restricted to the reversal phase, including both reversal training and the test based on reversed contingencies, it is likely that NA-OFC is involved in both creating new states and in the “online” use of the information included in this new state. Such a proposal is in accordance with popular theories of LC-NA system which suggest that a rise in NA activity allows for behavioral flexibility when a change in contingencies is detected (Aston-Jones et al 1997, Bouret & Sara 2005, Sadacca et al 2016).

## Conclusion

Our results confirm the involvement of ventral and lateral OFC in the updating and use of new action-outcome associations, and further demonstrate that NA input to the OFC is critical for these learning processes. Recent research has revealed a remarkable parcellation of cortical functions in goal-directed action (Parkes & Coutureau, 2019; Turner & Parkes, 2020). The current study provides a clear basis for an in-depth understanding of the cortical coordination involved in executive functions.

## DECLARATION OF INTERESTS

The authors declared that there is no actual or potential conflict of interest about this article.

## ACKNOWLEDGEMENTS

This work was supported by the French National Research Agency (CE37-0019 NORAD to E.C. and EJK) and by the Fondation pour la Recherche Médicale (FRM grant number ECO20160736024 to J-C.C.). Funding sources had no further role in study design, in the collection, analysis and interpretation of data, in the writing of the report and in the decision to submit the paper for publication. Microscopy was completed at the Bordeaux Imaging Centre, a service unit of CNRS-INSERM and Bordeaux University and member of the national infrastructure, France BioImaging. The authors thank Angélique Faugere (INCIA, CNRS, UMR 5287) for help with immunofluorescence and Yoan Salafranque (INCIA, CNRS, UMR 5287) for expert animal care. The authors are grateful to Alain Marchand for his help in every steps of this project.

## AUTHORS CONTRIBUTIONS

All authors had full access to all the data in the study and took responsibility for the integrity of the data and the accuracy of the data analysis. Conceptualization: E.C. & S.P.; Methodology: J-C C., E. K., M.L., M.W. & A.P.; Investigation: J-C. C., M.D. & A.P.; Formal Analysis: A.P., J-C C., E.C. & S.P.; Resources: E.C., E. K., M.W. & S.P.; Writing: S.P., A.P., J-C C., M.W. & E.C; Supervision: E.C., M.W. & S.P.; Funding EK & E.C.

## SUPPLEMENTAL FIGURES

**Suppl. Figure 1.**
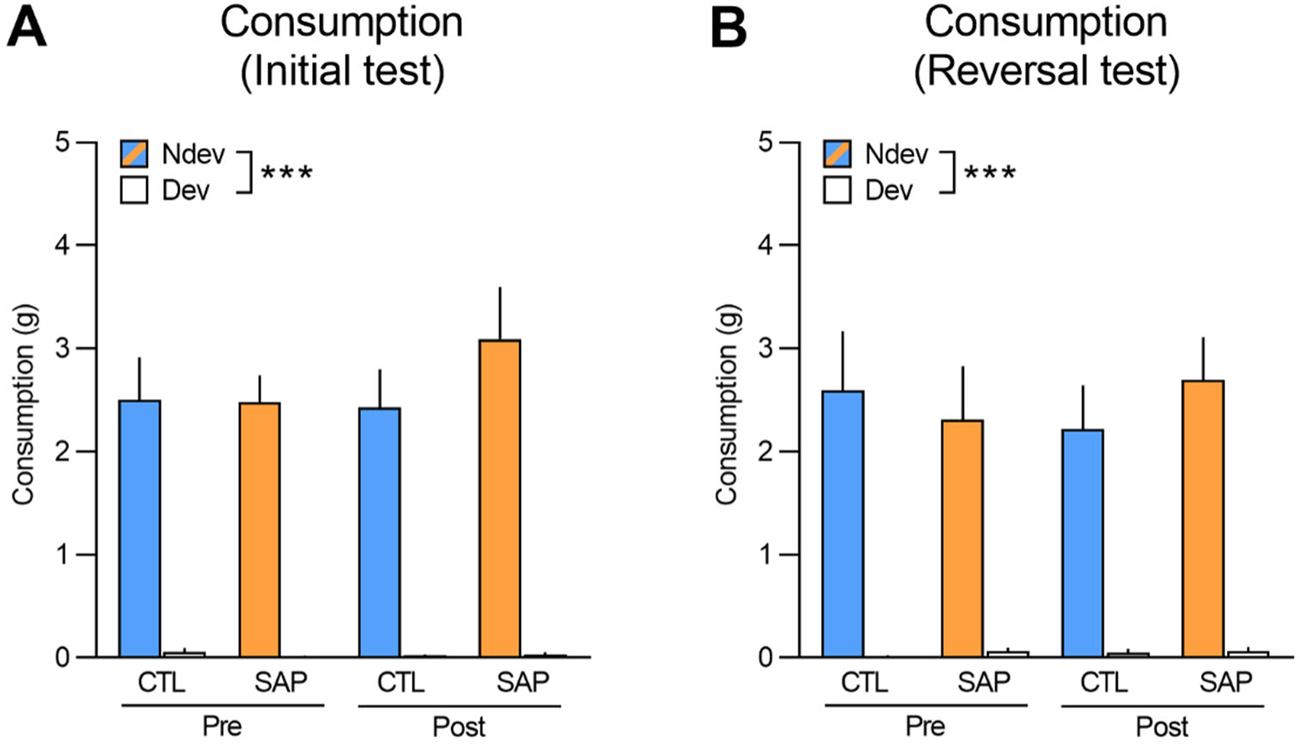
Consumption tests performed immediately after the initial (**A**) and reversal (B) instrumental tests in extinction. Rats were given access to both food rewards (10 g each) for 10 min. Statistics revealed a within-subjects effect of devaluation for both the initial (F_(1,53)_=168.94, p <0.001) and reversal tests (F_(1,53)_=97.25, p <0.001) and no main effects of group, treatment, or interactions between these factors (largest F_(1,53)_ value=0.57, p=0.45). Data are presented as mean + S.E.M. ***p <0.001.

**Suppl. Figure 2.**
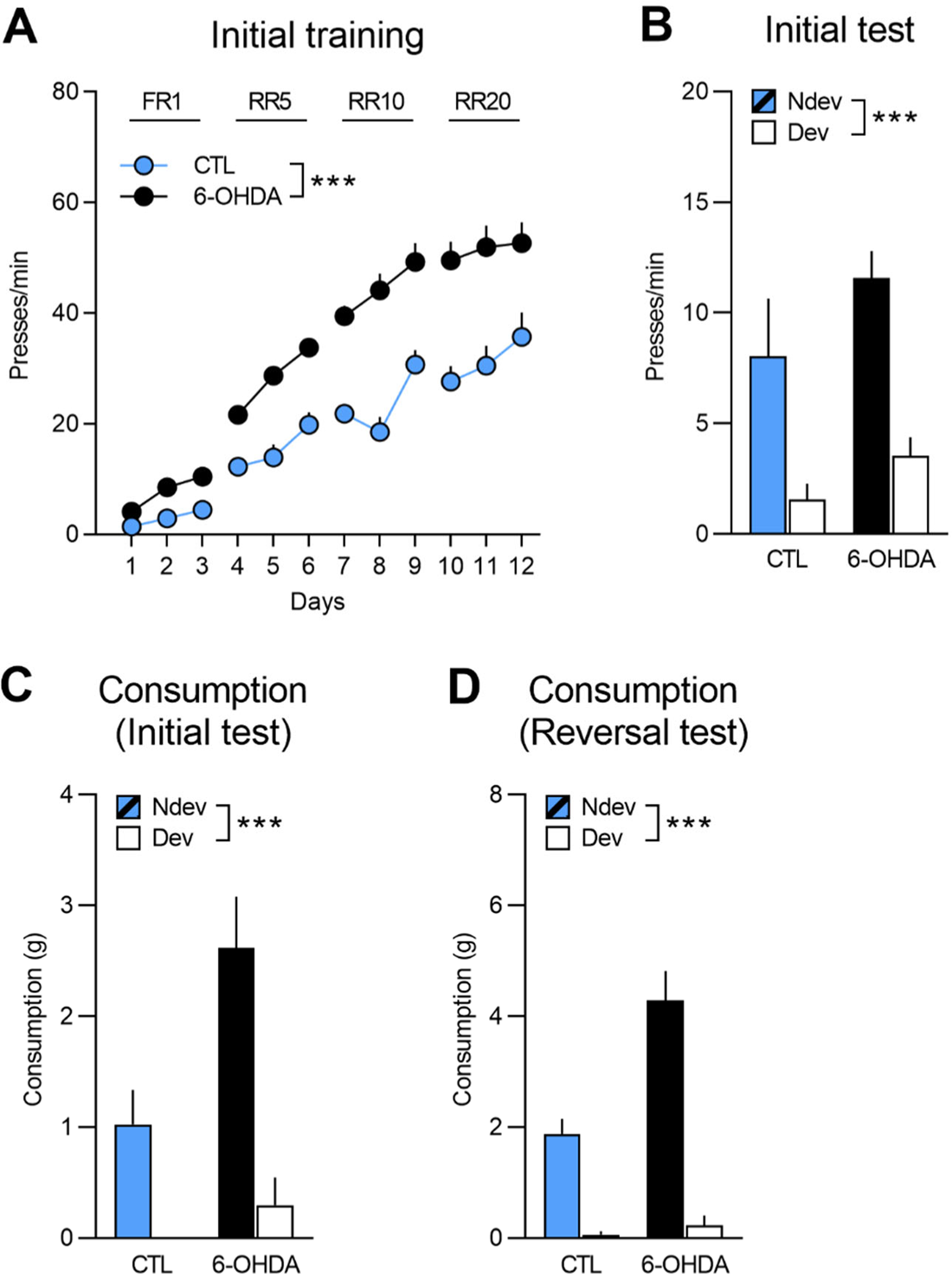
Initial training and test for rats to be injected with 6-OHDA (n = 12) and control rats (n = 8). (**A**) Training data is presented collapsed across the two actions (A1-O1; A2-O2). There was a main effect of group (F_(1,18)_=35.33, p <0.001), devaluation (F_(1,18)_=190.66, p <0.001) and group x devaluation interaction (F_(1,18)_=8.8, p <0.01). Simple effects confirmed that responding increased in both group 6-OHDA and group control (F_(1, 18)_=175.85, p <0.001 and F_(1,18)_=48.8, p <0.001, respectively). (**B**) Statistical analyses on the outcome devaluation test revealed an overall effect of devaluation (F_(1,18)_=39.37, p <0.001), no effect of group (F_(1,18)_=2.84, p=0.11), and no group x devaluation interaction (F_(1,18)_=0.46, p=0.51). (**C**) During the initial consumption test, both groups consumed more of the non-devalued outcome as indicated by a significant effect of devaluation (F_(1,18)_=18.69, p <0.001) and group (F_(1,18)_=9.6, p <0.01) but no group x devaluation interaction (F_(1,18)_=2.79, p=0.11). (**D**) The results from the reversal consumption test revealed a significant main effect of devaluation (F_(1,18)_=94.72, p <0.001) as well as a significant main effect of group (F_(1,18)_=10.0, p <0.01) and a significant group x devaluation interaction (F_(1,18)_=13.86, p <0.01), indicating that the difference between consumption of the non-devalued versus devalued food was actually greater for group 6-OHDA (devalued mean = 0.23; non-devalued mean = 4.29) than for group control (devalued mean = 0.06; non-devalued mean = 1.86). Nevertheless, simple effects analyses confirmed that both 6-OHDA and control groups consumed more of the non-devalued food than the devalued food (F_(1,18)_=15.05, p <0.01and F_(1,18)_=113.16, p <0.001, respectively). Data are presented as mean + S.E.M. ***p <0.001.

**Suppl. Figure 3.**
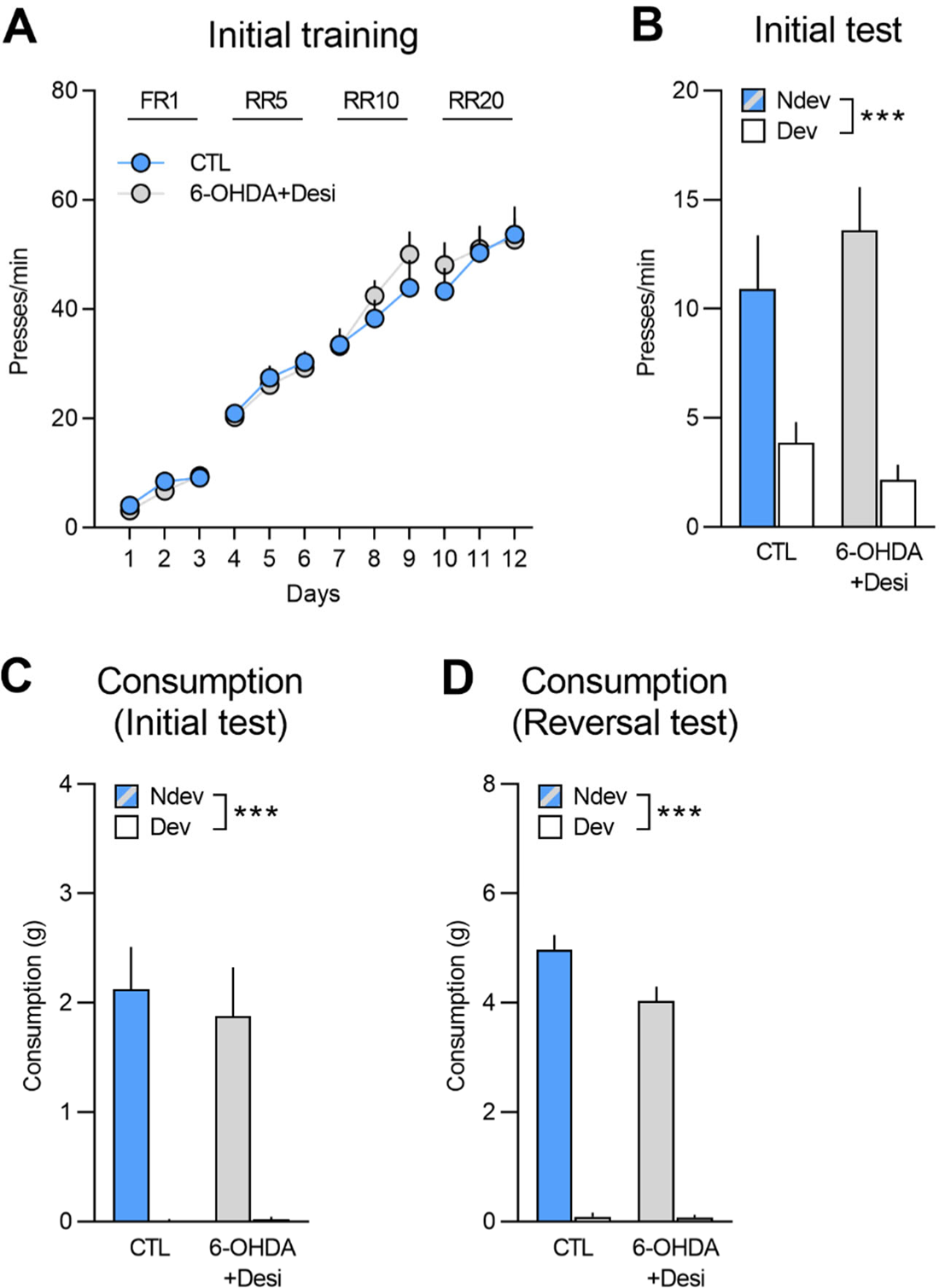
Initial training and test for rats to be injected with 6OHDA+Desi (n = 9) and control rats (n = 8). (**A**) Training data is presented collapsed across the two actions (A1-O1; A2-O2). There was a main effect of devaluation (F_(1,15)_=220.45, p <0.001) but no effect of group (F_(1,15)_=0.06, p=0.81) or group x devaluation interaction (F_(1,15)_=0.31, p=0.59). (**B**) Statistical analyses on the outcome devaluation test revealed an overall effect of devaluation (F_(1,15)_=44.36, p <0.001), no effect of group (F_(1,15)_=0.07, p=0.79), and no group x devaluation interaction (F_(1,15)_=2.5, p=0.13). (**C**) During the initial consumption test, both groups consumed more of the non-devalued outcome as indicated by a significant effect of devaluation (F_(1,15)_=45.57, p <0.001) but no effect of group (F_(1,15)_=0.15, p=0.70) or group x devaluation interaction (F_(1,15)_=0.19, p=0.67). (**D**) During the reversal consumption test both groups still consumed more of the non-devalued food (F_(1,15)_=466.43, p <0.001). Data are presented as mean + S.E.M. ***p <0.001.

**Suppl. Figure 4.**
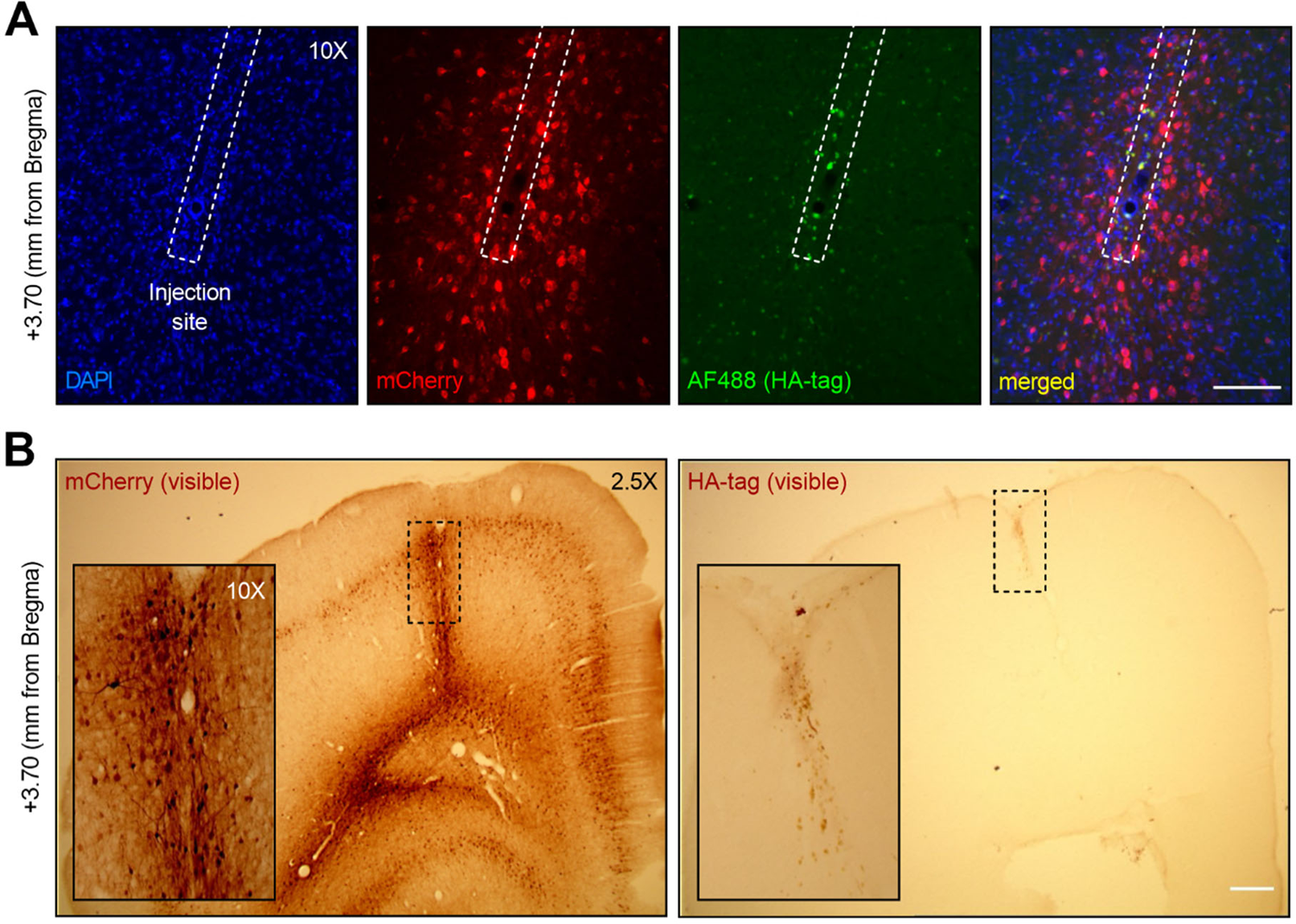
(**A**) Immunofluorescent staining for mCherry and HA (tag of inhibitory DREADDs) in one injection site of a representative rat injected with CAV2-PRS in the OFC. (**B**) DAB staining for mCherry and HA in one injection site of the same representative rat injected with CAV2-PRS in the OFC. In both cases, although mCherry-stained cell bodies can be seen all around the injection site (and above), there are no HA-stained cells. Scale bar (**A**): 100 μm; scale bar (**B**): 400 μm.

**Suppl. Figure 5.**
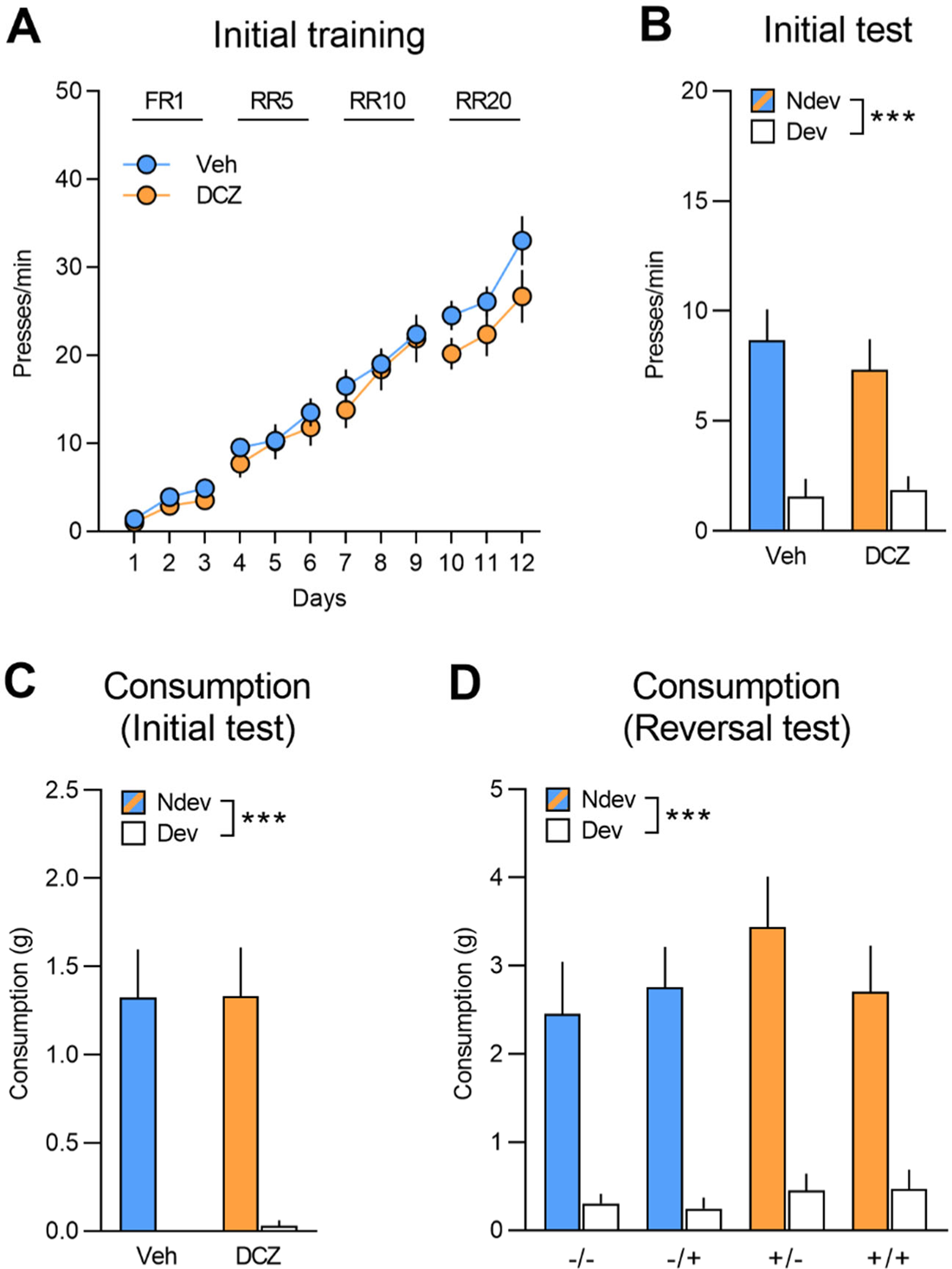
(**A**) Initial training for rats injected with CAV2-PRS in the OFC that would be allocated to group vehicle (Veh; n = 12) and group DCZ (n = 13) for the subsequent reversal training, data is presented collapsed across the two actions. Both groups acquired the instrumental response (F_(1,23)_=236.06, p <0.001) and there was no difference between groups (F_(1,23)_=0.53, p=0.47) or interaction (F_(1,23)_=0.45, p=0.51). (**B**) Initial instrumental test following satiety-induced devaluation. Statistics revealed a main effect of devaluation (F_(1,23)_=32.61, p <0.01) but no effect of group (F_(1,23)_=0.16, p=0.69) or a significant interaction between these factors (F_(1,23)_=0.30, p=0.59). (**C**, **D**) All groups consumed more of the non-devalued food than the devalued food during the consumption tests performed immediately after the initial (**C**) (F_(1,23)_=42.93, p <0.001) and reversal instrumental tests (**D**) (F_(1,23)_=74.19, p <0.001) with no main effects of group or treatment or interactions between these factors (largest F value=2.76, p=0.11). Data are presented as mean + S.E.M. ***p <0.001.

**Suppl. Figure 6.**
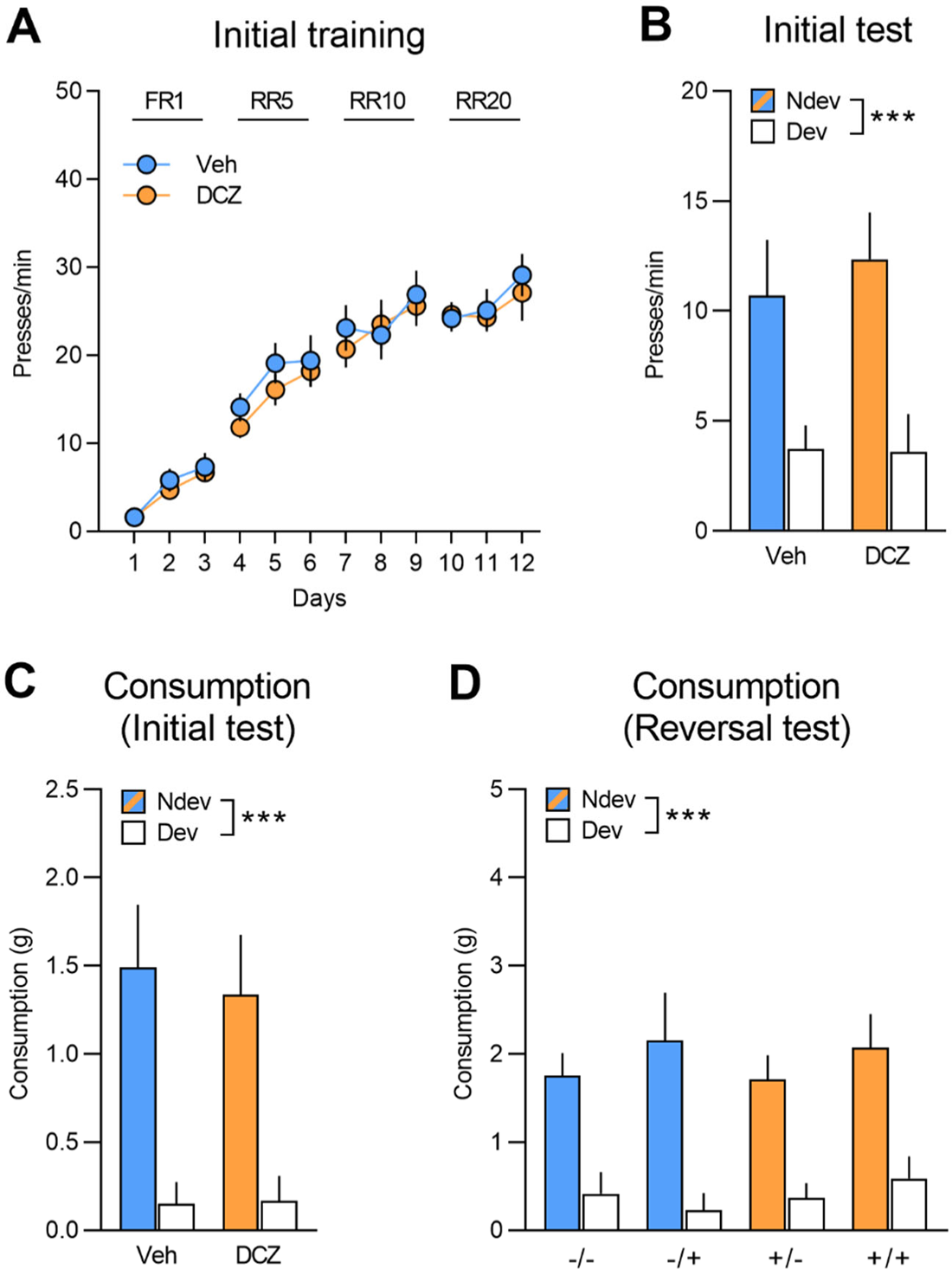
(**A**) Initial training for rats injected with CAV2-PRS in the mPFC that would be allocated to group vehicle (Veh n = 8) and group DCZ (n = 9) for the subsequent reversal training, data is presented collapsed across the two actions. Both groups acquired the instrumental response (F_(1,15)_=278.73, p <0.001) and there was no difference between groups (F_(1,15)_=1.37, p=0.26) or interaction (F_(1,15)_=0.16, p=0.70). (**B**) Initial instrumental test following satiety-induced devaluation. Statistics revealed a main effect of devaluation (F_(1,15)_=12.57, p=0.01) but no effect of group (F_(1,15)_=0.22, p=0.65) or a significant interaction between these factors (F_(1,15)_=0.16, p=0.70). (**C**, **D**) All groups consumed more of the non-devalued food than the devalued food during the consumption tests performed immediately after the initial (**C**) (F_(1,15)_=19.92, p <0.001) and reversal (**D**) instrumental tests (F_(1,15)_=51.18, p <0.001) with no main effects of group or treatment or interactions between these factors (largest F value=1.1, p=0.31). Data are presented as mean + S.E.M. ***p <0.001.

---

Detailed methods are provided in the online version of this paper and include the following:

## KEY RESOURCES TABLE

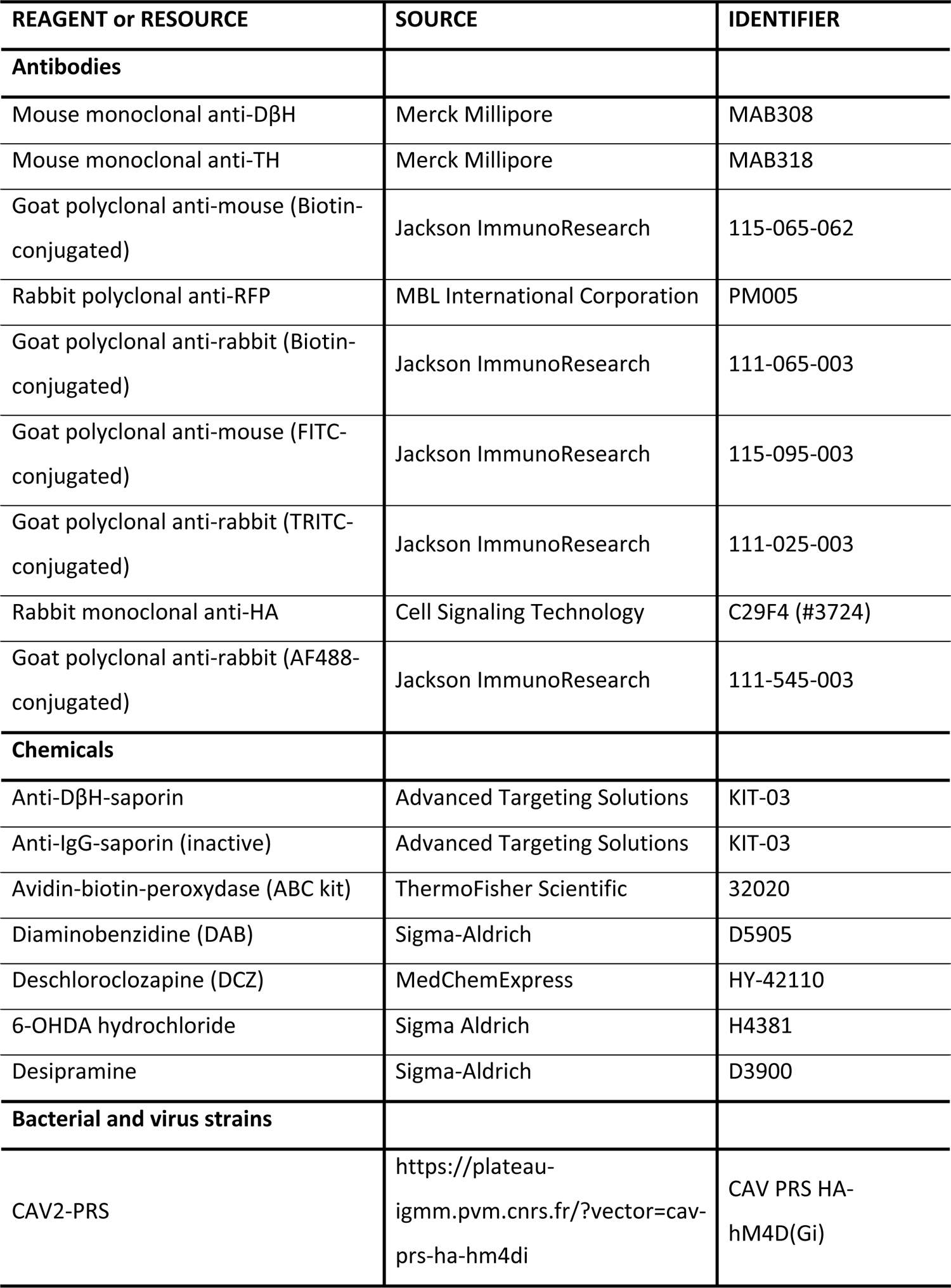

| REAGENT or RESOURCE | SOURCE | IDENTIFIER |
| --- | --- | --- |
| **Antibodies** |  |  |
| Mouse monoclonal anti-DβH | Merck Millipore | MAB308 |
| Mouse monoclonal anti-TH | Merck Millipore | MAB318 |
| Goat polyclonal anti-mouse (Biotin-conjugated) | Jackson ImmunoResearch | 115-065-062 |
| Rabbit polyclonal anti-RFP | MBL International Corporation | PM005 |
| Goat polyclonal anti-rabbit (Biotin-conjugated) | Jackson ImmunoResearch | 111-065-003 |
| Goat polyclonal anti-mouse (FITC-conjugated) | Jackson ImmunoResearch | 115-095-003 |
| Goat polyclonal anti-rabbit (TRITC-conjugated) | Jackson ImmunoResearch | 111-025-003 |
| Rabbit monoclonal anti-HA | Cell Signaling Technology | C29F4 (#3724) |
| Goat polyclonal anti-rabbit (AF488-conjugated) | Jackson ImmunoResearch | 111-545-003 |
| **Chemicals** |  |  |
| Anti-DβH-saporin | Advanced Targeting Solutions | KIT-03 |
| Anti-IgG-saporin (inactive) | Advanced Targeting Solutions | KIT-03 |
| Avidin-biotin-peroxydase (ABC kit) | ThermoFisher Scientific | 32020 |
| Diaminobenzidine (DAB) | Sigma-Aldrich | D5905 |
| Deschloroclozapine (DCZ) | MedChemExpress | HY-42110 |
| 6-OHDA hydrochloride | Sigma Aldrich | H4381 |
| Desipramine | Sigma-Aldrich | D3900 |
| **Bacterial and virus strains** |  |  |
| CAV2-PRS | https://plateau-igmm.pvm.cnrs.fr/?vector=cav-prs-ha-hm4di | CAV PRS HA-hM4D(Gi) |

## RESOURCE AVAILABILITY

### Lead contact

Further information and requests for resources should be directed to and will be fulfilled by the lead contact, Etienne Coutureau.

## EXPERIMENTAL MODEL AND SUBJECT DETAILS

### Animals and housing

A total of 136 male Long-Evans rats, aged 2-3 months, were obtained from the Centre d’Elevage Janvier (France). Rats were housed in pairs with *ad libitum* access to water and standard lab chow prior to behavioural experiments. Rats were handled daily for 3 days prior to the beginning of the experiments and were put on food restriction 2 days before behaviour to maintain them at approximately 90% of their *ad libitum* feeding weight. The facility was maintained at 21±1 °C on a 12 hr light/dark cycle (lights on at 8:00 am). Environmental enrichment was provided by tinted polycarbonate tubes and nesting material, in accordance with current French (Council directive 2013-118, February 1, 2013) and European (directive 2010-63, September 22, 2010, European Community) laws and policies regarding animal experimentation. The experiments received approval from the local Bordeaux Ethics Committee (CE50).

## METHODS DETAILS

### Stereotaxic surgery

For all experiments, rats were anesthetized with 5% inhalant isoflurane gas with oxygen and placed in a stereotaxic frame with atraumatic ear bars (Kopf Instruments) in a flat skull position. Anaesthesia was maintained with 1.5% isoflurane and complemented with a subcutaneous injection of ropivacaïne (a bolus of 0.1 mL at 2 mg/mL) at the incision site. After each injection, the injector was kept in place for an additional 10 min before being removed. Rats were given 4 weeks to recover following surgery. Injection sites were confirmed histologically after the completion of behavioural experiments.

In the first experiment (n = 57), we used a toxin selective for noradrenergic neurons (saporin; SAP) to target and deplete noradrenergic terminals in the OFC. For half of the rats (‘Pre’ groups, n = 29), surgery was performed before the initial instrumental training and testing phase, for the other half surgery was performed after the initial training and testing (‘Post’ groups, n = 28). Intracerebral injections were made using repeated pressure pulses delivered via a glass micropipette connected to a pressure injector (Picospritzer III, Parker). For SAP groups (Pre n = 15; Post n = 15), 0.1 µL of anti-DβH saporin (0.1 µg/µL) was bilaterally injected at one site targeting the OFC, while control (CTL) rats (Pre n = 14; Post n = 13) received 0.1 µL of inactive anti-IgG saporin (0.1 µg/µL). Injection coordinates (in mm from Bregma) were determined from the atlas of Paxinos and Watson (2014): +3.5 antero-posterior (AP), ±2.2 medio-lateral (ML) and −5.4 dorso-ventral (DV).

We then used a toxin selective for catecholaminergic neurons (6-OHDA hydrochloride) and a noradrenaline uptake-blocker (desipramine; Desi) to target and deplete dopaminergic neurons in the OFC. All rats underwent surgery after the initial instrumental training phase. Rats were then allocated to the full catecholaminergic depletion condition (group 6-OHDA n = 12; control n = 8) or the specific DA depletion condition (6OHDA+Desi n = 9; control n = 8). 6-OHDA (4 µg/µL) was dissolved in vehicle solution containing 0.9% NaCl and 0.1% ascorbic acid. A volume of 0.2 µL of 6-OHDA was bilaterally injected in the OFC at the same coordinates as for the first experiment. Animals in the CTL group received injections of the vehicle solution. Thirty minutes before the surgical procedure, animals in the 6-OHDA+Desi group received a systemic (i.p.) injection of desipramine (25 mg/mL) at a volume of 1 mL/kg.

In the chemogenetic experiments (n = 42), we employed a canine adenovirus type 2 (CAV2) vector equipped with a noradrenergic-selective synthetic promoter (PRS) and inhibitory DREADDs (hM4Di) to specifically target LC:OFC and LC:mPFC noradrenergic projections. This viral construct was designed and controlled at the Plateforme de Vectorologie de Montpellier, Institute of Molecular Genetics, Montpellier, https://plateau-igmm.pvm.cnrs.fr/?vector=cav-prs-ha-hm4di. All rats underwent surgery before the initial instrumental training phase. All animals received 1.0 μL bilateral injections of the adenovirus (titer 3.5 × 10^12 pp/mL), which were performed using a 10 μL Hamilton syringe and a stereotax-mounted injection pump (World Precision Instruments) at a flow rate of 100 nL/min. To target both ventral and lateral regions of the OFC, rats (n = 25) were injected at the following coordinates (mm from Bregma): +3.7 AP, ±2.0 ML, −5.0 DV and +3.2 AP, ±2.8 ML, −5.2 DV (2 injection sites per hemisphere). To target the mPFC, rats (n = 17) were injected at the following coordinates (mm from Bregma): +3.2 AP, ±0.6 ML, −3.6 DV (1 injection site per hemisphere).

### Behavioural apparatus

For all behavioural experiments, training and testing was conducted in 8 identical operant chambers (40 cm width x 30 cm depth x 35 cm height, Imetronic, Pessac, France) individually enclosed in sound and light resistant wooden chambers (74 x 46 x 50 cm). Each chamber was equipped with 2 pellet dispensers that delivered grain (Rodent Grain-Based Diet, 45mg, Bio-Serv) or sugar (LabTab Sucrose Tablet, 45 mg, TestDiet) pellets into a food port when activated. For instrumental conditioning, two retractable levers were located on each side of the food port. Each chamber had a ventilation fan producing a background noise of 55 dB. During the session, the chamber was illuminated by four LEDs in the ceiling. Experimental events were controlled and recorded by a computer located in the room and equipped with the POLY software (Imetronic).

### Behavioural protocol

#### Initial training and test

The training procedure was adapted from Parkes et al (2018). On days 1-3, rats were trained to retrieve food pellets from the food port. During each daily session, 40 sugar and 40 grain pellets were delivered pseudo-randomly every 60 s, on average. Following food port training, rats received 12 daily sessions of instrumental training, during which they were required to learn initial action-outcome (A-O) associations. During these sessions, each lever, in alternation, was presented twice for a maximum of 10 min or until 20 outcomes were earned. The inter-trial interval between lever presentations was 2.5 min (i.e., the session could last up to 50 min and the rats could obtain a maximum of 80 food pellets). The A-O associations and the order of lever presentations were counterbalanced between rats and days. During the first three sessions, lever pressing was continuously reinforced with a fixed ratio (FR) 1 schedule. Then, the probability of receiving an outcome was reduced, first with a random ratio (RR) 5 schedule (days 4-6), then with an RR10 (days 7-9,) and an RR20 schedule (days 10-12).

Outcome devaluation tests were performed one day after the last instrumental training session. First, to induce sensory-specific satiety (Rolls 1986), rats received access to one of the two outcomes (20 g) for 1 hr in a set of plastic feeding cages to which they were previously habituated. Immediately after the satiety procedure, rats were returned to the operant chambers where they were given a choice test in extinction (i.e. unrewarded) with both levers available for 10 min. The devalued (sated) food was counterbalanced between rats. Following the extinction test, animals were returned to the plastic feeding cages and given a consumption test of satiety-induced devaluation, during which they received 10 min concurrent access to both types of food pellets (10 g of each). The amount consumed of each pellet type was measured to confirm that the satiety-induced devaluation was effective and that rats were able to distinguish between the sensory features of the different food pellets.

#### Reversal training and test

Following the initial phase, rats were trained on reversed A-O associations with a procedure adapted from previous studies from our laboratory (Fresno et al 2019, Parkes et al 2018). Specifically, the identity of outcomes was switched so that rats had to update previously established A-O associations, always keeping a RR20 schedule of reinforcement. Following reversal training, outcome devaluation tests were conducted in the same manner as previously described.

### Chemogenetics

The DREADD agonist deschloroclozapine (DCZ) was dissolved in dimethyl sulfoxide (DMSO) to a final volume of 50 mg/mL, aliquoted in small tubes (50 μL) and stored at −80°C (stock solution). For behavioural experiments, our stock solution was diluted in physiological saline to a final injectable volume of 0.1 mg/kg and administered systemically (i.p.) 40-45 min prior to testing at a volume of 10 mL/kg. Fresh injectable solutions were prepared from stock aliquots on the day of the usage. DCZ was always handled in dim light conditions.

### Histology

At the end of all behavioral experiments, rats were injected with a lethal dose of sodium pentobarbital (Exagon® Euthasol) and perfused transcardially with 60 mL of saline followed by 260 mL of 4% paraformaldehyde (PFA) in 0.1 M phosphate buffer (PB). Brains were removed and post-fixed in the same PFA 4% solution overnight and then transferred to a 0.1 M PB solution or to a 0.1 M PB with 30% saccharose solution (6-OHDA experiment). Subsequently, 40 µm coronal sections were cut using a VT1200S Vibratome (Leica Microsystems) or freezing microtome for the 6-OHDA experiment. Every fourth section was collected to form a series. DAB staining was performed for DβH (for the saporin and the 6-OHDA experiments), TH (6-OHDA experiment), HA and mCherry (chemogenetic experiments).

Free-floating sections were first rinsed (4 x 5 min) in 0.1 M phosphate buffer saline (PBS) containing 0.3% Triton X-100 (PBST) and then incubated in PBST containing 0.5% (for mCherry) or 1% (for DβH and TH) hydrogen peroxide solution (H2O2) for 30 min in the dark. Further rinses (4 x 5 min) in PBST and a 1 hr incubation in blocking solution (PBST containing 3% goat serum) followed. Sections were then incubated with the primary antibody (mouse monoclonal anti-DBH, 1/10000; mouse monoclonal anti-TH, 1/2000; rabbit monoclonal anti-HA, 1/1000; rabbit polyclonal anti-RFP, 1/2000) diluted in blocking solution for 24 hr (for mCherry) or 48 hr (for DβH and TH) at 4°C. After rinses (4 x 5 min) in PBS (for DβH and TH) or PBST (for mCherry), sections were placed in a bath containing the secondary antibody (biotinylated goat anti-mouse, 1/1000; biotinylated goat anti-rabbit, 1/1000) diluted in PBS (for DβH and TH) or PBST containing 1% goat serum (for mCherry) for 2 hr at room temperature. Following rinses (4 x 5 min) in PBS (for DβH and TH) or PBST (for mCherry), they were then incubated with the avidin-biotin-peroxydase complex (1/200 in PBS for DβH and TH; 1/500 in PBST for mCherry) for 90 min in the dark at room temperature. H_2_O_2_ was added to the solution before the final staining with DAB was made (10 mg tablet dissolved in 50 mL of 0.1 M Tris buffer). Stained sections were finally rinsed with 0.05 M Tris buffer (2 x 5 min) and 0.05 M PB (2 x 5 min), before being collected on gelatin-coated slides using 0.05 M PB, dehydrated (with xylene for DβH and TH), mounted and cover-slipped using the Eukitt mounting medium.

Immunofluorescence staining was also performed for DβH, mCherry and HA (chemogenetic experiments). Free-floating sections were first rinsed in PBS (4 x 5 min) and PBST (3 x 5 min), before being incubated in blocking solution for 1 hr at room temperature. Sections were then incubated with the primary antibody (mouse monoclonal anti-DβH, 1/1000; rabbit polyclonal anti-RFP, 1/1000; rabbit monoclonal anti-HA, 1/1000) diluted in blocking solution for 24 hr at 4°C. Following rinses in PBS (4 x 5 min), they were then incubated with the secondary antibody (goat polyclonal anti-mouse FITC-conjugated, 1/400; goat polyclonal anti-rabbit TRITC-conjugated, 1/200; with goat polyclonal anti-rabbit, 1/1000) diluted in PBS for 2 hr in the dark at room temperature. Stained sections were finally rinsed with PSB (4 x 5 min), before being collected on gelatin-coated slides using 0.05 M PB, dehydrated, mounted and cover-slipped using Fluoroshield with DAPI mounting medium.

## QUANTIFICATION AND STATISTICAL ANALYSIS

### Fiber loss

To measure fiber density in the saporin and 6-OHDA experiments, we used the protocol described in Cerpa et al (2019). We examined sections at +4.4, +3.7 and +3.0 (mm from Bregma) using a Nanozoomer slide scanner with a 20X lens (Hamamatsu Photonics). Digital microphotographs of regions of interest (ROI, square windows of 300 x 300 µm, 1320 x 1320 pixels) in each hemisphere were examined under a 20X virtual lens with the NDP.view 2 freeware (Hamamatsu Photonics). Each ROI was outlined according to Paxinos and Watson (2014). Quantification of DβH- and TH-positive fibers was performed using an automated method developed in the laboratory with the ImageJ software (Cerpa et al 2019). Specifically, a digitized version of the microphotograph was converted to black and white by combining blue, red and green channels (weights 1, −0.25 and −0.25), subjected to a median filter (radius 3 pixels) in order to improve the signal-to-noise ratio, smoothed with a Gaussian filter (radius 8), and subtracted from the previous picture to isolate high spatial frequencies. Large stains were further eliminated by detecting them in a copy of the image. The picture was then subjected to a fixed threshold (grey level 11) to extract stained elements, and the relative volume occupied by fibers estimated by the proportion of detected pixels in the ROI. As a control for poor focus, the same images were analysed a second time while allowing lower spatial frequencies (Gaussian filter radius 20). The ratio between the proportions of pixels detected by the two methods was used as a criterion to eliminate blurry images.

### Experimental design and data analysis

Each rat was assigned a unique identification number that was used to conduct blind testing and statistical analyses. Behavioural data and fibre volume were analysed using sets of between and within orthogonal contrasts controlling the per contrast error rate at alpha = 0.05 (Hays, 1963). Simple effects analyses were conducted to establish the source of significant interactions. Statistical analyses were performed using PSY Statistical Program (http://www.psy.unsw.edu.au/research/research-tools/psy-statistical-program; Kevin Bird, Dusan Hadzi-Pavlovic, and Andrew Issac © School of Psychology, University of New South Wales) and graphs were created using GraphPad Prism.

All experiments employed a between-x within-subjects behavioural design. In the first experiment, the between-subject factors were group (Pre versus Post) and treatment (control versus saporin) and the within-subject factors were training day (acquisition data) or devaluation (responding on lever associated with non-devalued or devalued outcome) for the test data. In the 6-OHDA experiments, the between-subject factor was group (control vs 6-OHDA or control vs 6OHDA+Desi) and the within-subject factor was training day (acquisition data) or devaluation (test data). To analyse DβH and TH fibre volume, the between-subject factor was group (experiment 1: control versus saporin; experiment 2: control, 6OHDA+Desi, or 6OHDA) and the within-subject factor was region (VO versus LO) for the OFC. There was no within-subject factor for the quantification of fibres in the mPFC. In the final chemogenetics experiment, the between-subject factor was treatment during reversal acquisition (vehicle versus DCZ) and the within-subject factors were training day (acquisition data) or treatment during reversal test (vehicle versus DCZ) and devaluation (test data).

## Notes

### Competing Interest Statement

The authors have declared no competing interest.

